# Local and network-level dynamics of alpha and beta rhythms during visuospatial attention in autism

**DOI:** 10.64898/2026.11.18.767297

**Authors:** Leonie Polzer, Sara Boxhoorn, Magdalena Schütz, Andreas M. Grossmann, Hannah Mössinger, Nico Bast, Christina Luckhardt, Christine M. Freitag

**Author notes:** corresponding author Correspondence to Leonie Polzer Department of Child and Adolescent Psychiatry, Psychosomatic Medicine and Psychotherapy, University Hospital Frankfurt, Goethe-University, Deutschordenstraße 50, 60528 Frankfurt am Main, Germany.

## Abstract

Visuospatial attention is altered in autism, yet the underlying neural mechanisms remain incompletely understood. Neural oscillatory activity coordinates distributed processes supporting attentional selection and their flexible adjustment to changing environmental demands. Altered oscillatory organization may therefore provide a mechanistic basis for atypical attentional processing in autism. We tested this theory using EEG recordings from 80 autistic and neurotypical children and youth performing a spatial cueing task with valid and invalid cues. We quantified alpha- and beta-band measures of local power and inter-trial phase clustering (ITPC) together with power- and phase-based measures of posterior-frontal long-range connectivity. Across validity conditions, autistic participants showed a more frontal alpha distribution, reduced posterior alpha lateralization and posterior beta ITPC, and higher frontal beta power. Crucially, autistic participants also presented reduced neural adaptation across attentional conditions, reflected in attenuated validity-dependent modulation of alpha power and of posterior-frontal beta connectivity, with elevated beta connectivity particularly in the valid cue condition. This neural pattern was accompanied by greater reaction-time slowing in the invalid compared with the valid cue condition. Within autistic participants, beta dynamics showed nominal associations with restricted and repetitive behaviors that did not survive multiple-comparison correction. Together, these findings suggest that visuospatial attention in autism is characterized by altered local oscillatory organization, combined with reduced modulation of local and distributed dynamics as attentional demands change. Alterations in alpha and beta activity may reflect reduced spatially selective attentional gating alongside greater maintenance of established attentional states.

## Introduction

The ability to flexibly allocate and reallocate attention in response to changing behavioral demands is crucial for navigating everyday environments. Efficient visuospatial attention relies on using prior information, such as predictive cues, to guide attentional orienting towards currently relevant locations, while remaining able to disengage and redirect attention when behaviorally relevant stimuli appear elsewhere. Visuospatial attention is altered in autism spectrum disorder (ASD), as part of a broader profile of attentional characteristics that may affect development and adaptive functioning [1, 2]. Behavioral assessments indicate slower cue-guided orienting across the lifespan [3]. Some studies additionally reported difficulties in disengaging and reorienting attention outside prioritized locations, although these findings have been less consistent [3–6]. Thus, visuospatial attention in autism may involve less efficient cue-guided orienting, and a reduced flexibility in adjusting established attentional priorities. However, the neural mechanisms underlying these potentially distinct alterations remain insufficiently characterized.

Visuospatial attention relies on coordinated activity within visual and fronto-parietal networks that support selective orienting and reorienting to behaviorally relevant events [7–9]. Its neural implementation involves both local activity and interactions across distributed regions, which are thought to rely on coordinated oscillatory activity [10, 11]. Both local and network-level dynamics can be characterized through complementary measures of oscillatory amplitude and phase consistency, reflecting the strength and temporal organization of rhythmic neural activity, respectively.

At the local level, power in the alpha (8–12 Hz) and beta (13–30 Hz) frequency bands is implicated in visuospatial attention. Alpha power is spatially modulated by attention, typically decreasing contralateral to attended locations and increasing over task-irrelevant regions [12–14]. These modulations have been proposed to reflect changes in cortical excitability and attentional gating [15, 16]. Beta power has been linked to top-down control of sensorimotor and cognitive states, with increases associated with maintaining ongoing states [17, 18], and content-specific beta activity accompanying the reactivation of endogeneous representations [19].

Beyond oscillatory power, phase consistency provides a complementary measure of local oscillatory dynamics, reflecting the temporal organization of neural response timing. Inter-trial phase clustering (ITPC) quantifies the consistency of oscillatory phase across repeated events and is modulated in visual and parieto-frontal regions during attention [20, 21].

At the network level, attention requires coordinated interactions across distributed cortical regions, which can be characterized through power-based and phase-based connectivity, reflecting interregional covariation of oscillatory power and phase synchronization, respectively [22]. Increased long-range alpha phase-based connectivity has been implicated in the coordination of spatially distributed processing during visuospatial attention [14, 16, 23, 24], while beta connectivity supports top-down interactions between frontal and visual regions, with evidence from both phase- and power-based measures [25–27]. Together, these local and network-level oscillatory mechanisms provide complementary mechanisms through which attentional processing is implemented.

In autism, differences in oscillatory activity have been reported across both resting-state and task contexts, with growing evidence that their expression depends on current processing demands. At rest, alterations in alpha activity are among the more consistent findings across development, whereas differences in beta activity are less robust [28–30]. During task performance, autistic individuals have shown attenuated modulation of alpha activity during visual and multisensory processing, including reduced suppression to relevant input and reduced enhancement over task-irrelevant regions [31, 32]. Findings on beta activity likewise suggest atypical task-related dynamics in autism, with both increased and decreased activity reported, alongside attenuated modulation between experimental conditions [33–35]. Reduced ITPC across frequencies encompassing alpha and beta bands further suggests less temporally consistent stimulus-related responses in autism [33, 36].

At the network level, both increased and decreased long-range functional connectivity in lower and mid-range frequency bands have been reported in autism, with findings varying across tasks, and connectivity measures, although reductions have been reported more frequently [37–39]. In some studies, task-related alpha and beta connectivity have been found to be increased during visual discrimination, auditory and picture-naming tasks [40–42], suggesting that connectivity alterations depend on the cognitive context in which networks are engaged. Findings of attenuated modulation of alpha and beta phase-based connectivity during response inhibition [43] additionally point towards reduced adjustment of neural connectivity to changing demands. These findings raise the possibility that the underlying neural mechanisms of attentional alterations in autism may comprise both altered oscillatory organization and altered reconfiguration of local and distributed neural dynamics as attentional demands change.

Together, previous findings point to several candidate mechanisms of oscillatory activity that may contribute to attentional differences in autism, but direct evidence from studies using controlled visuospatial attention paradigms remains limited. It therefore remains unclear whether differences are primarily expressed in local response magnitude or temporal consistency, in distributed network coordination, or across multiple levels simultaneously. Moreover, little is known about whether such differences are present when prior information can be used to guide attention, when attentional priorities need to be adjusted, or during both. Examining oscillatory dynamics across these conditions could therefore clarify both the neural mechanisms of visuospatial attention in autism and its flexibility as attentional demands change.

Against this background, we investigated whether adaptive visuospatial attention in autistic compared with neurotypical children and youth differs in local neural processing, distributed network coordination, and their reconfiguration across changing attentional demands. Using a spatial cueing paradigm, we contrasted validly cued targets, for which prior information could be used to establish a spatial priority, with invalidly cued targets requiring disengagement and reorienting. Local processing was assessed through oscillatory power and ITPC, and network-level processing through power-based and phase-based connectivity, focusing on alpha and beta activity because of their complementary roles in spatial selection and the maintenance of neural states. We expected attenuated spatial alpha power modulation and altered long-range connectivity in autism, while directional hypotheses for beta activity and ITPC were not specified. Alongside differences in overall oscillatory organization, a further focus of our analysis was to examine whether neural measures were differentially modulated by cue validity, testing for altered reconfiguration when established attentional priorities had to be changed. Secondary analyses examined relations among neural measures, associations with behavioral performance, and clinical symptoms.

## Materials and methods

### Participants and phenotypic measures

The study included 43 autistic and 37 age-matched neurotypical participants aged 10-17 years (Table 1), with IQ ≥ 70 assessed using standardized composite scores from verbal comprehension and perceptual organization subscales of German Wechsler intelligence scales [44, 45]. Expert clinical autism diagnoses were confirmed using the Autism Diagnostic Observation Schedule (ADOS/ADOS-2 [46, 47]) and Autism Diagnostic Interview-Revised (ADI-R; [48]). Inclusion required an ADOS/ADOS-2 classification of ‘autism’ when ADI-R data were unavailable (n = 2). Autism symptom severity was characterized using ADOS/ADOS-2 calibrated severity scores for social affect and restricted/repetitive behavior (RRB), and inattention using the inattention subscale of the ADHD parent rating scale [49]. Neurotypical participants were excluded for elevated psychopathology (internalizing, externalizing, or total T-score ≥ 60 in the Child Behaviour Checklist/4-18R (CBCL/4-18R; [50]). Additional exclusion criteria are described elsewhere [51]. Written informed consent was obtained from participants and caregivers. The study was approved by the ethics committee of the Medical Faculty, University Hospital Frankfurt (416/17).

**Table 1.** Sample description.

|  | **ASD**<br>**(n=43)** | **NT**<br>**(n=37)** | **p-value** |
|---|---|---|---|
| Sex (n male/n female) | 40/3 | 27/10 | .034 |
| Age (years) | 14.52 (2.20)<br>[10.50-17.83] | 15.30 (1.68)<br>[11.33-17.75] | .076 |
| IQ | 100.12 (15.52)<br>[70-130] | 106.30 (9.49)<br>[88-128] | .032 |
| Handedness (LQ) | 60.61 (46.60)<br>[-80-100] | 73.64 (41.26)<br>[-100-100] | .189 |
| CBCL internalizing (T-score) | 63.02 (8.43)<br>[50-85] | 46.74 (7.88)<br>[36-58] | < .001 |
| CBCL externalizing (T-score) | 56.77 (11.72)<br>[36-81] | 42.56 (6.58)<br>[36-56] | < .001 |
| CBCL total (T-score) | 63.60 (10.52)<br>[45-90] | 42.97 (7.21)<br>[31-57] | < .001 |
| ADOS/ADOS-2 CSS | 6.55 (1.80)<br>[4-10] | - | - |
| ADI-R social interaction | 15.88 (5.01)<br>[4-27] | - | - |
| ADI-R communication | 12.63 (5.44)<br>[1-24] | - | - |
| ADI-R rituals | 4.83 (2.29)<br>[0-10] | - | - |

*Note:* Values are presented as mean (SD) [range], unless otherwise specified. P-values are derived from two-sided unpaired t-tests; sex differences were assessed using Pearson's chi-squared test. ADOS/ADOS-2 CSS could not be calculated for one ASD participant, as their age exceeded the standardized range provided in the manual. ADI-R values were missing for two participants. ASD, autism spectrum disorder; NT, neurotypical participants; IQ, intelligence quotient; CBCL, Child Behaviour Checklist/4-18R; ADOS, Autism Diagnostic Observation Schedule; CSS, calibrated severity score; ADI-R, Autism Diagnostic Interview-Revised.

### Visuospatial cueing task

Figure 1 summarizes the experimental paradigm and analytical workflow. Participants performed a Posner spatial cueing task (Figure 1A) with two boxes flanking a central fixation cross. Following a variable inter-trial interval (500-1100 ms), one box was highlighted as a spatial cue (50 ms), followed by a target (500 ms) at a stimulus onset asynchrony (SOA) of 100 or 250 ms. The cue validly or invalidly indicated target location. Participants indicated target location by left or right keypress as quickly and accurately as possible. Of 512 cue-target trials, 70% were valid (n = 356) and 30% invalid (n = 156). Additional cue-only and target-only trials (n = 64 each) were excluded from analyses.

**Figure 1.**
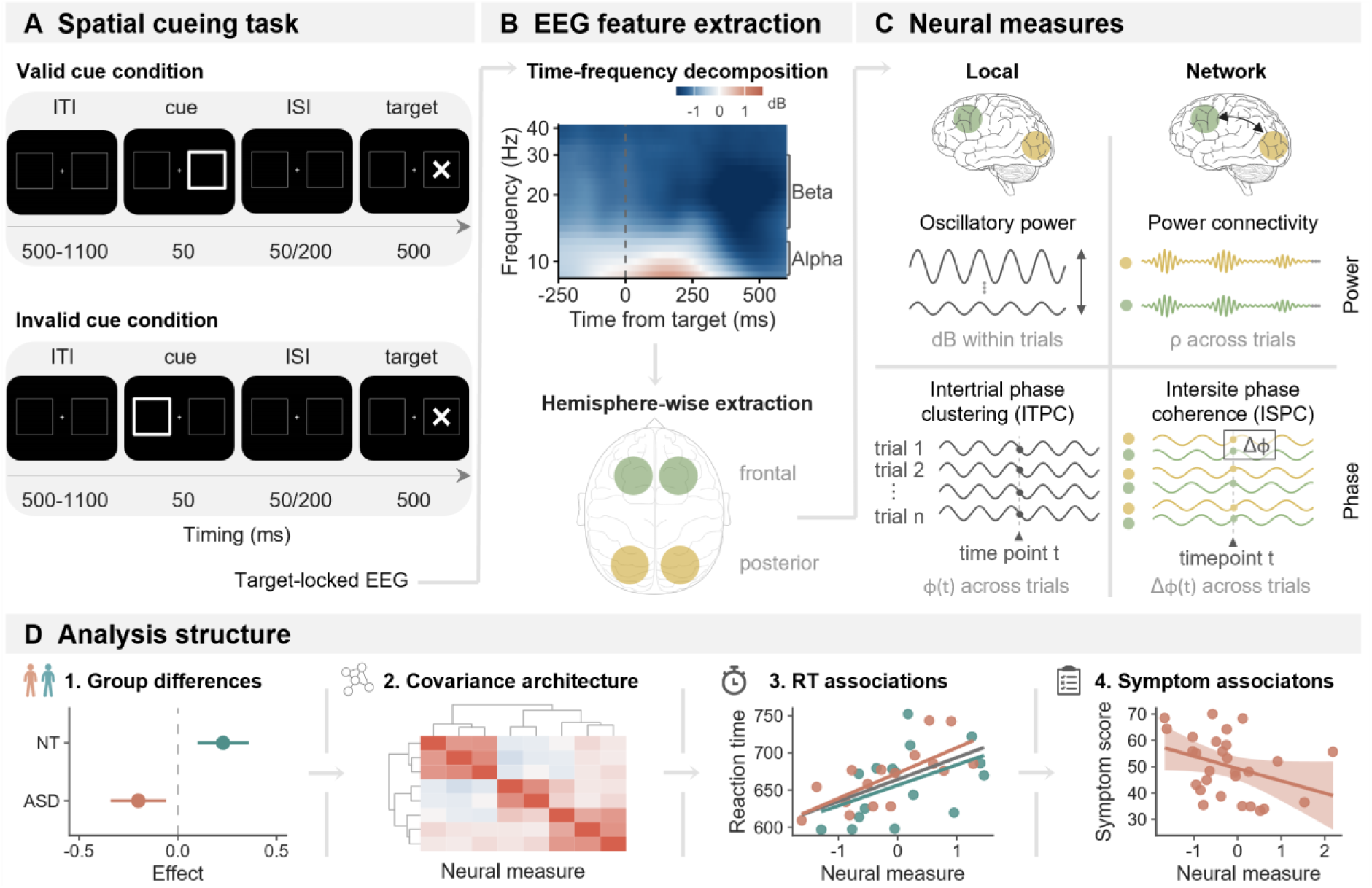
Overview of the methodological framework. (A) Valid and invalid cue conditions of the Posner spatial cueing task. (B) Target-locked time-frequency decomposition and hemisphere-wise extraction of frontal and posterior activity. The time-frequency map shows an illustrative grand-average pooled across groups, validity conditions, hemispheres, and regions. (C) Local measures comprised oscillatory power and inter-trial phase clustering (ITPC), whereas network measures comprised power connectivity and inter-site phase coherence (ISPC); Ф denotes phase at a given time point and ΔФ the posterior-frontal phase difference. (D) Analyses assessed group differences, covariance architecture, reaction-time associations, and associations with ASD symptom severity.

### EEG acquisition and preprocessing

EEG was recorded using 64 Ag/AgCl electrodes positioned according to the extended international 10-20 system, with FCz as recording reference and an additional electrode for vertical EOG (∼1 cm below right eye). Impedances were maintained below 10 kΩ. Data were acquired at 1000 Hz using BrainAmp MR-Plus amplifiers and BrainVision Recorder software (online anti-aliasing bandpass filter: 0.01-250 Hz, 12 dB/octave).

EEG data were preprocessed using EEGLab (v2025.0.0; [52]) and custom MATLAB (R2025a) scripts. Data were downsampled to 500 Hz, cleaned of line noise (50 Hz, 100 Hz), and screened for bad channels using Clean Rawdata (Flatline ≥ 5 s, correlation < 0.65, or line-noise > 5 SD) and visual inspection. Data were average-referenced, and high-pass filtered at 0.1 Hz. Extended infomax ICA was performed on a 1 Hz high-pass filtered data set after manual removal of segments with gross artifacts. Artifactual components (eye movements, muscle activity, line noise, ECG) were identified using ICLabel [53] and visual inspection, and removed from the 0.1 Hz filtered dataset. Rejected channels were subsequently interpolated using spherical splines, followed by average re-referencing.

Target-locked epochs (−1.25 to +1.3 s) were extracted, providing padding for time-frequency analyses. Automated screening identified potentially artifactual epochs (peak-to-peak amplitude > 200 µV or robust z > 6.0; voltage jumps > 80 µV; flatness < 0.5 µV (SD); or high-frequency power ratio (40-80 Hz / 8-20 Hz) > 3.0 with robust z > 3.5), which were manually reviewed before rejection.

Trials with reaction times < 200 ms (ASD: 3.22%; NT: 2.79%), > 1000 ms or omitted responses (ASD: 3.01%, NT: 2.22%), and trials with incorrect responses (ASD: 2.98%, NT: 2.41%) were excluded. For multiple responses within the valid window, only the first was evaluated. The proportion of retained trials did not differ between groups (valid trials: ASD = 76.02%, NT = 80.50%, p = .119; invalid trials: ASD = 74.97%, NT = 80.23%, p = .071).

#### Time-frequency decomposition and regions of interest

Time-frequency (TF) decomposition was performed using complex Morlet wavelets implemented in FieldTrip [54] with FFT-based convolution applied to the full epochs before cropping to the analysis interval to reduce edge artifacts (30 logarithmically spaced frequencies, 8-60 Hz; logarithmically increasing number of 4-9 wavelet cycles). The TF grid was cropped to −500 to +1000 ms relative to target onset. Posterior (PO7, PO8, PO3, PO4, O1, O2), and frontal (FC3, FC4, FC1, FC2, F1, F2) regions of interest (ROIs) were defined based on visual inspection of grand averages collapsed across groups and validity conditions. Electrode clusters were classified as ipsilateral or contralateral to target location. Frequency bands comprised alpha (8-12 Hz) and beta (14-29 Hz) ranges. Custom MATLAB scripts quantified local oscillatory dynamics using total power and inter-trial phase clustering (ITPC), and network-level interactions using across-trial power-based and phase-based connectivity. Power was analyzed ipsi- and contralaterally, whereas ITPC and connectivity analyses focused on the contralateral hemisphere (Figure 1B-C).

#### Local dynamics

Total power was calculated as the trial-wise squared magnitude of the complex analytic signal and expressed in decibels (dB) relative to a trial-specific baseline (10 × log₁₀[post/baseline]). The baseline extended from −500 to −200 ms relative to cue onset to avoid contamination by cue-locked responses. Baseline normalization was performed separately for each electrode, frequency, and trial before averaging across electrodes and frequencies within each band, and subsequently across the 0-500 ms post-target interval. As a secondary measure of hemispheric asymmetry, power lateralization was calculated as ipsilateral minus contralateral baseline-normalized power.

Inter-trial phase clustering (ITPC) quantified stimulus-locked phase consistency across trials as the vector length of normalized complex phase angles. ITPC was calculated at each electrode, frequency and time point, and averaged across electrodes within each ROI, frequencies within bands, and the 0-500 ms post-target interval. To account for trial-count dependence, ITPC was averaged across 100 iterations of random subsampling of 20 trials without replacement within each validity condition × target location × SOA condition. One neurotypical participant with fewer than 20 artifact-free invalid trials was excluded from phase-based analyses.

#### Network-level interactions

Power-based connectivity was quantified as trial-to-trial co-modulation of oscillatory power between posterior and frontal ROIs. Spearman rank correlations across trials quantified baseline-normalized power co-modulation over 0-500 ms post-target for each participant, validity condition, hemisphere, and frequency band. Correlation coefficients were subsequently Fisher z-transformed.

Phase-based connectivity was quantified using inter-site phase coherence (ISPC, also termed phase-locking value [PLV]), a measure of phase synchronization reflecting the consistency of interregional phase differences across trials at corresponding time-frequency points. ISPC was calculated at each frequency and time point for individual electrode pairs connecting ROIs, and subsequently averaged across pairs, frequencies within bands, and the 0-500 ms post-target interval. Trial counts were matched using the same subsampling procedure as for ITPC.

### Statistical analyses

Statistical analyses were performed in R (v4.4.2, [55]; Figure 1D). All regression models adjusted for age, IQ, and sex, with participant included as a random intercept in linear mixed-effects models (LMMs). Multiple comparisons were controlled using the Benjamini-Hochberg false discovery rate (FDR) correction, with analysis-specific correction families. Standardized model coefficients (β) are reported as effect-size estimates. Significant interactions were decomposed using FDR-adjusted post hoc contrasts, reported as standardized estimated marginal mean differences (Δβ) with unadjusted 95% confidence intervals.

Reaction time (RT) and local oscillatory power were analyzed at the trial level, whereas ITPC and network-level measures used participant-level estimates, with group, validity, and their interaction as fixed effects in LMMs. Local models additionally included ROI and its interactions with group and validity. Separate models were fitted for alpha and beta bands and, for power, for ipsilateral and contralateral hemispheres. Omnibus effects were evaluated using Type III tests with Satterthwaite-approximated degrees of freedom. FDR correction was applied to each omnibus effect within neural measures across frequency bands and, for power, hemispheres. Results focus on significant group-related effects; complete omnibus tests, additional ROI × validity contrasts, and simulation-based power analyses are provided in Supplementary Tables S1-S6, Supplementary Information S1-S3, and S4, respectively.

To characterize multivariate neural organization beyond individual measures, participant-level Pearson correlations were calculated across ROI-specific and, for power, hemisphere-specific neural measures, separately by group and validity. Correlations used pairwise-complete observations and were FDR-corrected across the 120 unique pairs within each group and validity condition. Matrix correspondence was quantified by correlating corresponding coefficients (r_matrix_), and differences between matrices were summarized using their root-mean-square difference (RMSE). Group differences in condition-specific correlation matrices, overall correlation strength, and validity-dependent changes in matrix structure were assessed using 50,000 permutations of diagnostic group membership, preserving paired condition data where applicable.

Neural-behavioral associations were examined using mixed-effects models with standardized RT as the outcome. Power and ITPC models simultaneously included posterior and frontal measures and their respective interactions with validity, whereas connectivity models included the respective connectivity measure, validity, and their interaction. Models were fitted separately by frequency band and, for power, hemispheres. Corresponding effects were FDR-corrected across models within each neural measure. FDR-significant neural effects were followed up in exploratory group-moderation models using group × neural-measure interactions or group × validity × neural-measure interactions for validity-dependent effects.

Exploratory analyses within the autistic group examined associations between neural alterations identified in the primary analyses and clinical symptoms in social affect, RRB, and inattention. Multivariable linear regressions examined validity condition-averaged neural activity and valid-minus-invalid modulation for each phenotype, with one representative marker from each implicated neural domain entered simultaneously. Continuous variables were standardized within complete-case samples, with HC3 heteroscedasticity-consistent standard errors and 95% confidence intervals. FDR correction was applied across the three phenotypes separately for each neural marker and score type.

## Results

### Behavioral performance

Mean RTs for valid and invalid cue conditions were 402.20 ms (SD = 130.23) and 441.42 ms (SD = 117.02) in autistic participants, and 384.19 ms (SD = 112.38) and 419.94 ms (SD = 98.8) in neurotypical participants. Both groups showed a RT cost of invalid relative to valid cues, as indicated by faster responses in the valid than invalid condition (ASD: Δβ = −0.515, 95% CI [−0.544, −0.486], p_adj_ < .001; NT: Δβ = −0.455, 95% CI [−0.485, −0.425], p_adj_ < .001). This RT cost was greater in autistic than neurotypical participants (group × validity: F(1, 31,820) = 8.03, p = .005, Δβ = 0.060, 95% CI [0.019, 0.101]). Groups did not differ within validity conditions (both p_adj_ ≥ .497).

### Oscillatory dynamics

#### Local band power

Time-frequency representations generally indicated increased alpha and decreased beta power relative to baseline (Supplementary Figure S1). Power showed ROI-dependent validity effects across frequency bands and hemispheres (all p_adj_ ≤ .029). The valid compared to the invalid condition was characterized by higher ipsilateral posterior but lower contralateral power, with larger contralateral reductions in posterior than frontal regions. Secondary analyses showed greater posterior than frontal power lateralization during the valid condition across frequency bands. During the invalid condition, this regional pattern was absent for alpha and shifted towards greater frontal than posterior lateralization for beta (see Supplementary Information S1-S2).

Validity-dependent modulation of ipsilateral alpha power differed between groups (group × validity: F(1,63720.72) = 7.95, p_adj_ = .019; Figure 2A). Neurotypical participants showed higher ipsilateral alpha power in the valid than invalid condition (Δβ = 0.074, 95% CI [0.050, 0.098], p_adj_ <. 001), whereas this effect did not survive FDR correction in autistic participants (Δβ = 0.026, 95% CI [0.003, 0.049], p_adj_ = .054, between-group difference: Δβ = −0.048, 95% CI [−0.081, −0.015]).

**Figure 2.**
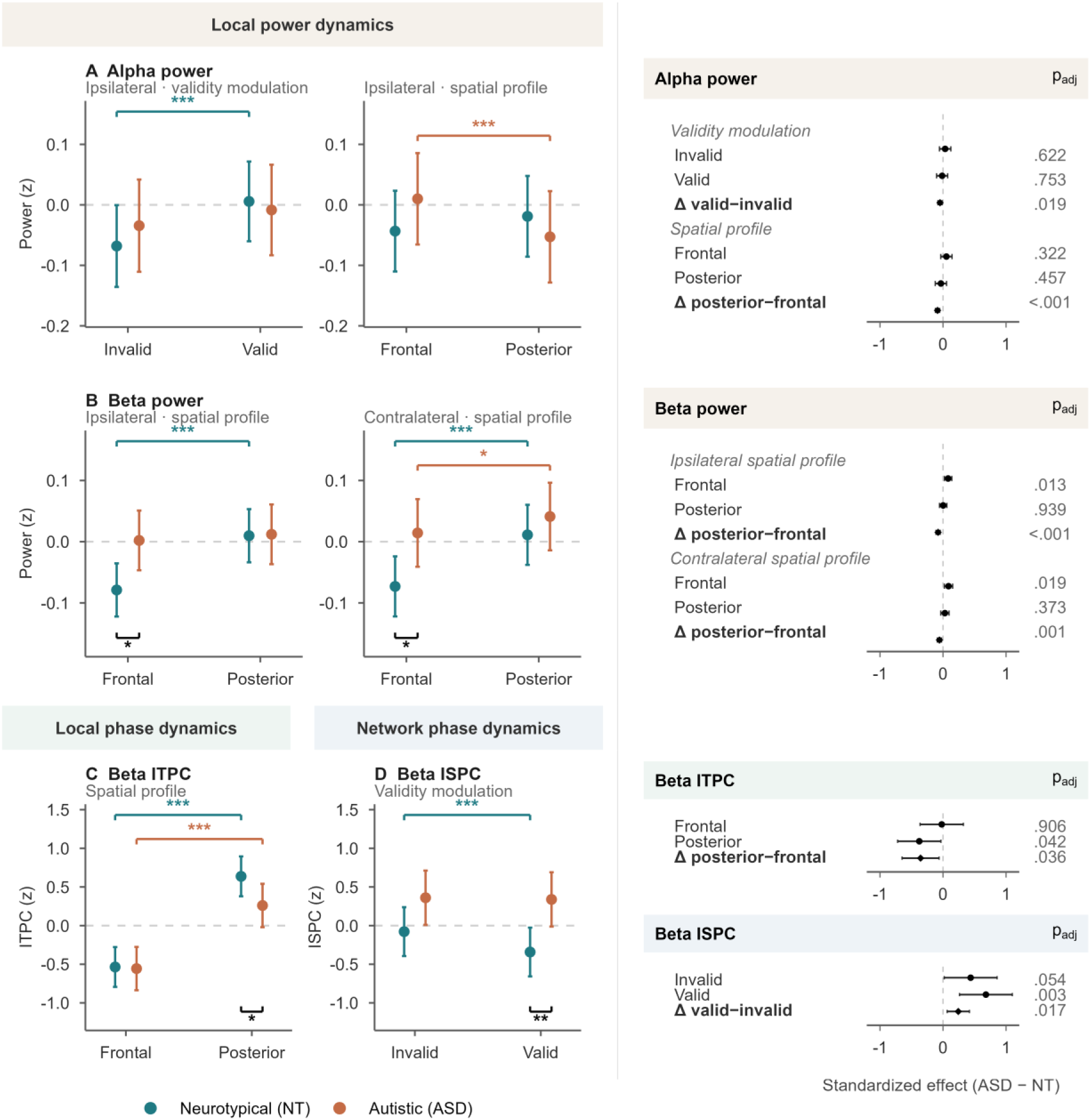
Group differences in local and network oscillatory dynamics. Summary of the neural measures showing FDR-corrected effects involving group, together with the corresponding post-hoc contrasts in: (A) Alpha power, (B) beta power, (C) beta inter-trial phase clustering (ITPC), and (D) beta inter-site phase coherence (ISPC). Dot-and-whisker plots show estimated marginal means and 95% confidence intervals for neurotypical (NT) and autistic (ASD) participants. Coloured brackets indicate within-group contrasts across validity or region of interest, whereas black brackets indicate between-group effects; only post-hoc contrasts surviving Benjamini-Hochberg FDR correction are annotated (* p_adj_ < .05, ** p_adj_ < .01, *** p_adj_ < .001). Forest plots on the right show the corresponding standardized ASD−NT group contrasts (circles) and group interaction effects (diamonds) with 95% confidence intervals; positive values indicate larger values or modulation in the ASD group. Confidence intervals reflect estimates from trial-level models for power and participant-level models for ITPC and ISPC. Δ valid−invalid and Δ posterior−frontal represent between-group differences in validity modulation and spatial profile, respectively. ITPC, inter-trial phase clustering; ISPC, inter-site phase coherence; FDR, false discovery rate.

Across validity conditions, spatial power distribution also differed between groups. Ipsilateral alpha power showed a group × ROI interaction (F(1, 63,716.05) = 26.55, p_adj_ < .001; Figure 2A), qualified by a lower posterior than frontal alpha power in autistic participants (Δβ = −0.063, 95% CI [−0.086, −0.040], p_adj_ < .001), whereas this regional difference was not significant in neurotypical participants (Δβ = 0.024, 95% CI [0.0005, 0.048], p_adj_ = .091; between-group difference: Δβ = −0.087, 95% CI [−0.121, −0.054]). Secondary lateralization analyses showed greater posterior alpha lateralization in neurotypical than autistic participants (Δβ = 0.059, 95% CI [0.021, 0.096], p_adj_ = .003; see Supplementary Information S2).

Beta power similarly showed bilateral group-by-ROI interactions (ipsilateral: F(1, 63,715.84) = 21.11, p_adj_ < .001, contralateral: F(1, 63,715.90) = 11.29, p_adj_ = .001; Figure 2B), with higher frontal beta power in autistic than neurotypical participants (ipsilateral: Δβ = 0.081, 95% CI [0.023, 0.139], p_adj_ = .013; contralateral: Δβ = 0.087, 95% CI [0.022, 0.153], p_adj_ = .019), and no posterior group differences (both p_adj_ ≥ .373). Additionally, neurotypical participants showed higher posterior than frontal beta power in both hemipsheres (ipsilateral: Δβ = 0.089, 95% CI [0.064, 0.113], p_adj_ < .001; contralateral: Δβ = 0.084, 95% CI [0.060, 0.108], p_adj_ < .001). This regional differentiation was non-significant for the ipsilateral hemisphere and smaller for the contralateral hemisphere in autistic participants (ipsilateral: Δβ = 0.010, 95% CI [−0.013, 0.033], p_adj_ = .532, between-group difference: Δβ = −0.079, 95% CI [−0.112, −0.045]; contralateral: Δβ = 0.027, 95% CI [0.004, 0.050], p_adj_ = .031, between-group difference: Δβ = −0.057, 95% CI [−0.091, −0.024]).

#### Inter-trial phase clustering (ITPC)

Alpha ITPC showed no significant effects of group, validity, or their interactions with ROI (all p_adj_ ≥ .415). Beta ITPC showed an ROI × validity interaction (F(1, 232.37) = 5.65, p_adj_ = .037), with greater posterior than frontal ITPC and lower frontal ITPC in the valid than invalid condition (see Supplementary Information S3).

Additionally, beta ITPC showed a group × ROI interaction (F(1, 232.37) = 5.66, p_adj_ = .036; Figure 2C). Neurotypical participants exhibited higher posterior beta ITPC than autistic participants (Δβ = 0.377, 95% CI [0.034, 0.720], p_adj_ = .042), with no group differences in the frontal ROI (p_adj_ = .906). Both groups showed greater posterior than frontal ITPC (NT: Δβ = 1.173, 95% CI [0.957, 1.389], p_adj_ < .001; ASD: Δβ = 0.816, 95% CI [0.617, 1.015], p_adj_ < .001), with a smaller difference in autistic compared to neurotypical participants (between-group difference: Δβ = −0.356, 95% CI [−0.650, −0.063]).

#### Network-level dynamics

Posterior-frontal power-based connectivity showed no group, validity, or interaction effects (all p_adj_ ≥ .270). Alpha phase-based connectivity (ISPC) was lower in the valid than invalid condition (F(1, 77.28) = 5.60, p_adj_ = .020, Δβ = −0.134, 95% CI [−0.245, −0.023]), with no additional effects of group.

Beta ISPC showed a group × validity interaction (F(1, 77.07) = 7.25, p_adj_ = .017; Figure 2D). Autistic participants exhibited higher beta ISPC than neurotypical participants in the valid condition (Δβ = 0.680, 95% CI [0.262, 1.099], p_adj_ = .003), whereas the difference in the invalid condition did not survive FDR correction (Δβ = 0.438, 95% CI [0.019, 0.858], p_adj_ = .054). Additionally, neurotypical participants showed higher beta ISPC in the invalid than valid condition (Δβ = 0.264, 95% CI [0.134, 0.394], p_adj_ < .001), whereas autistic participants showed no validity effect (Δβ = 0.022, 95% CI [−0.097, 0.141], p_adj_ = .721, between-group difference: Δβ = 0.242, 95% CI [0.066, 0.418], p_adj_ = .017). Group-related findings remained qualitatively unchanged after excluding participants with elevated ADHD symptoms (Tables S7-S11).

#### Covariation of local and network-level measures

Exploratory analyses of correlations among local and network-level neural measures revealed broadly similar covariation patterns between autistic and neurotypical participants in both validity conditions (valid: r_matrix_ = .771, invalid: r_matrix_ = .865; Figure 3), without significant whole-pattern group differences (valid: RMSE = .245, pperm = .191; invalid: RMSE = .171, p_perm_ = .901), and no differences in overall correlation strength or direction (all p_perm_ ≥ .280). However, neural covariation patterns differed more strongly between valid and invalid conditions in neurotypical than autistic participants (within-group RMSE = .193 versus .124; ΔRMSE = .069, p_perm_ = .045), and a trend towards greater group differences in valid than invalid conditions (ΔRMSE =.074, p_perm_=.054).

**Figure 3.**
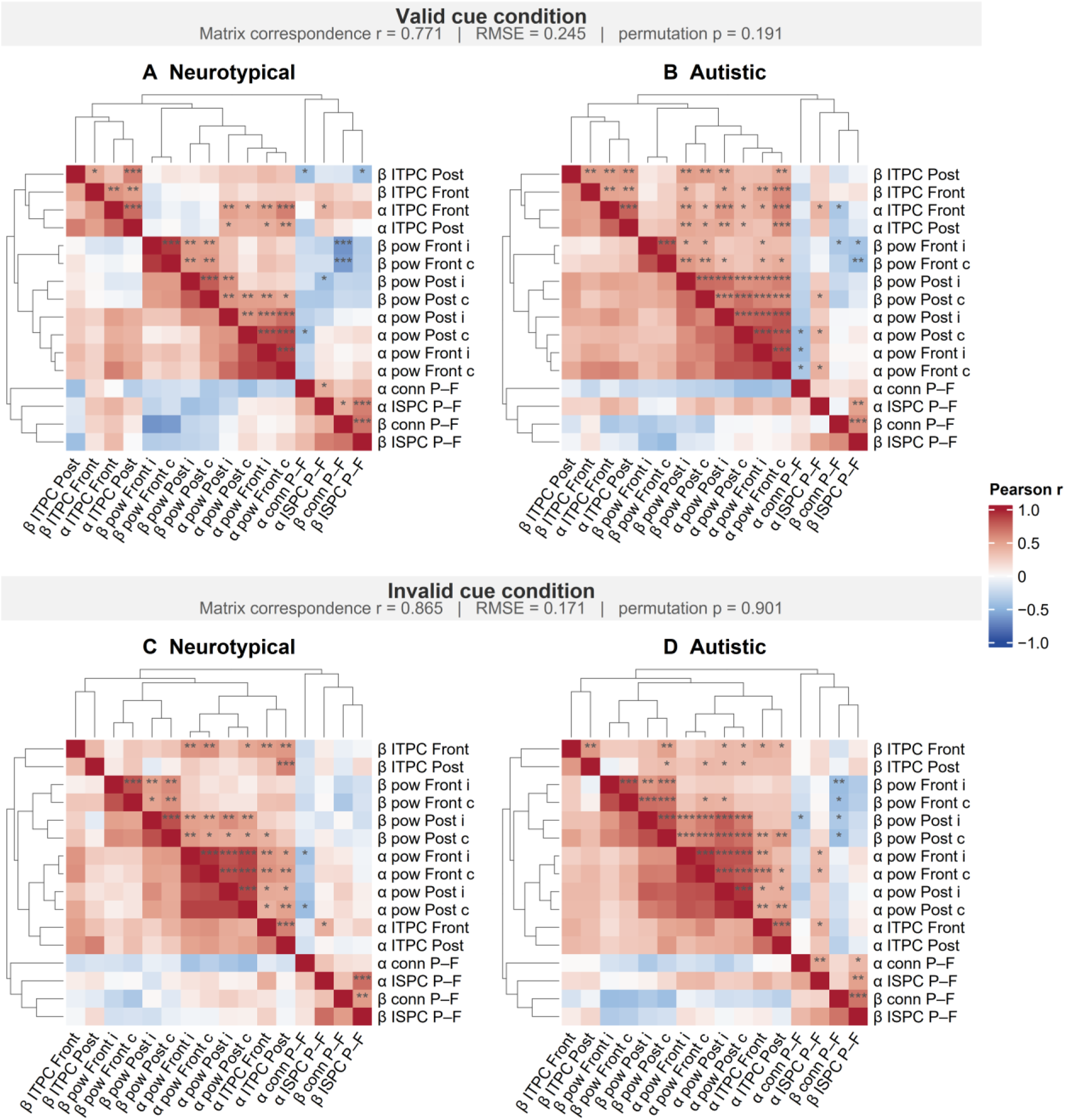
Interrelations among local and network-level neural measures. Group-wise pearson correlation matrices are shown separately for neural activity during the valid condition (A, B) and the invalid condition (C, D). Within each condition, a full-sample common hierarchical order was derived from pooled signed correlations using average-linkage clustering and applied to both group matrices. Dendrogram branches depict rank-scaled heights for visualization while preserving the original clustering structure and variable order. Asterisks denote correlations surviving within-group Benjamini-Hochberg FDR correction across all unique correlations: *p_adj_ < .05, **p_adj_ < .01, and ***p_adj_ < .001. Matrix correspondence represents the correlation between corresponding group-specific coefficients; RMSE summarizes their overall numerical discrepancy, with permutation *p*-values based on 5,000 random reassignments of group membership. Axis labels indicate frequency band (α, alpha; β, beta), neural measure (pow, power; ITPC, inter-trial phase clustering; conn, power connectivity; ISPC, inter-site phase coherence), region (Front, frontal; Post, posterior), hemisphere (i, ipsilateral; c, contralateral), and connectivity pair (P–F, posterior-frontal).

### Associations between neural dynamics and behavior

Higher ipsilateral posterior alpha power was associated with faster RT in the invalid (power x validity: F(1, 31,820.05) = 9.89, p_adj_ = .007; Δβ = −0.040, 95% CI [−0.058, −0.021], p_adj_ < .001), but not in the valid condition (Δβ = −0.004, 95% CI [−0.016, 0.008], p_adj_ = .526). Higher posterior beta power in both hemispheres was also associated with faster RT independent of validity (contralateral: F(1, 31,824.49) = 4.65, p_adj_ = .042; Δβ = −0.012, 95% CI [−0.023, −0.001]; ipsilateral: F(1, 31,824.59) = 6.81, p_adj_ = .018; Δβ = −0.015, 95% CI [−0.026, −0.004]). No frontal-power effect survived FDR correction. Greater posterior alpha lateralization (F(1, 31,821.57) = 10.22, p_adj_ = .003; Δβ = −0.018, 95% CI [−0.028, −0.007]), and frontal beta lateralization (F(1, 31,817.50) = 8.48, p_adj_ = .007; Δβ = −0.016, 95% CI [−0.026, −0.005]) were also associated with faster RT. Moreover, higher contralateral frontal alpha ITPC was associated with faster RT (F(1, 92.59) = 6.41, p_adj_ = .026, Δβ = −0.123, 95% CI [−0.219, −0.028]). Neither connectivity measure was significantly associated with RT (all p_adj_ ≥ .165). Full models are provided in Tables S12-S16. Exploratory analyses provided no evidence that the identified local neural-behavioral associations differed between groups.

### Associations between altered neural markers and clinical symptoms

In autistic participants, associations between observed autism symptoms, parent-rated inattention, and one representative marker per altered neural domain were examined: region-averaged ipsilateral alpha power, frontal beta power, posterior beta ITPC, and posterior-frontal beta ISPC (Figure 4). No associations emerged with ADOS/ADOS-2 social-affect severity or parent-rated inattention (all raw p ≥ .154). At the uncorrected level, greater ADOS/ADOS-2 RRB severity was associated with higher frontal beta power (β = .392, 95% CI [.052, .732], p = .025), higher posterior-frontal beta ISPC (β = .235, 95% CI [.008, .461], p = .043), and greater posterior beta ITPC in the invalid relative to the valid condition (β = −.331, 95% CI [−.648, −.013], p = .042). However, none of these effects survived FDR correction (p_adj_ ≥ .076).

**Figure 4.**
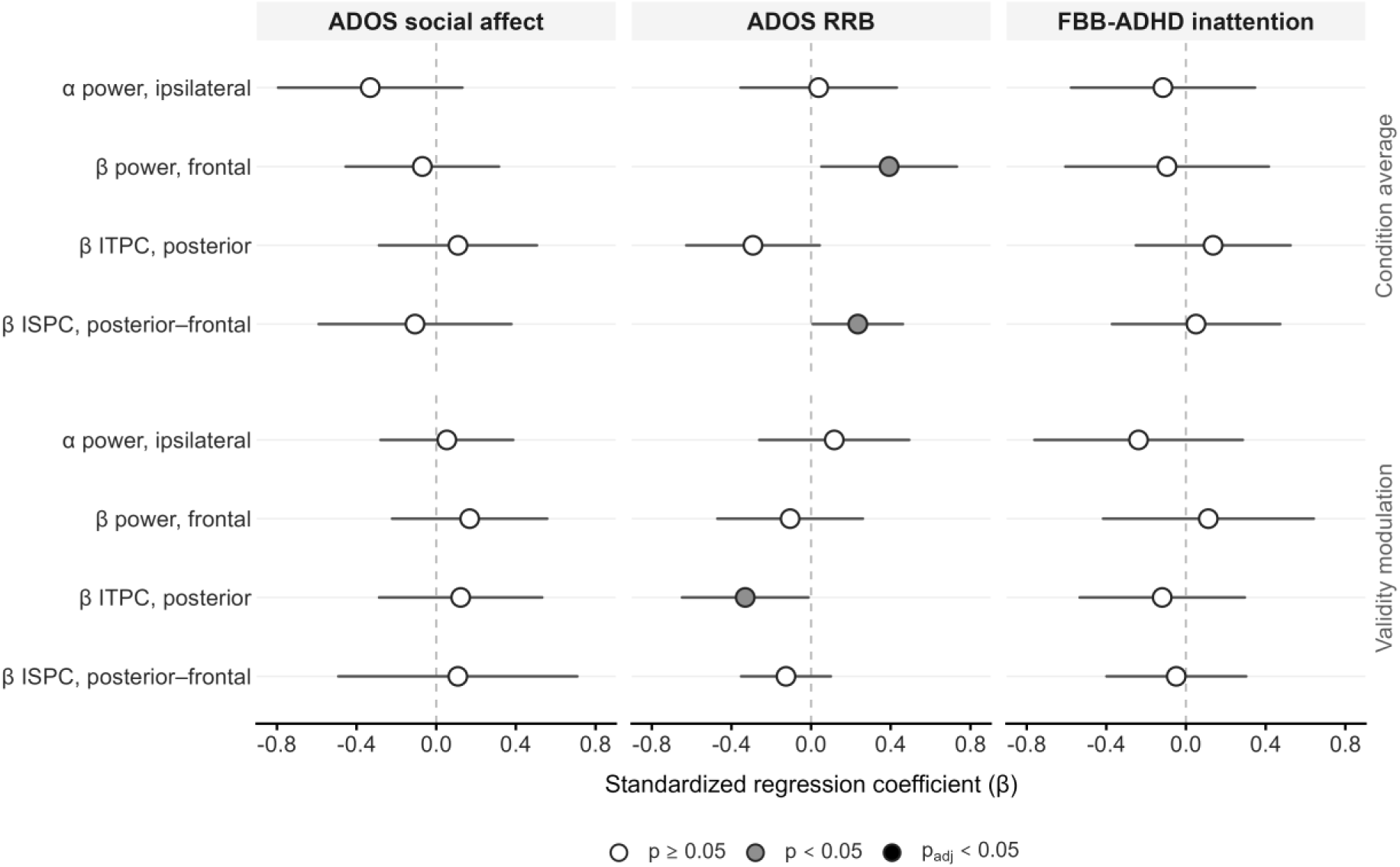
Associations between neural markers and symptom dimensions within the autistic group. Points show standardized coefficients from multivariable models adjusted for the other neural markers, age, IQ, and sex; horizontal lines indicate HC3 robust 95% confidence intervals. Upper and lower rows show condition-averaged activity and validity modulation (valid-minus-invalid), respectively. Open, grey, and black circles indicate p ≥ .05, raw p < .05, and p_adj_ < .05. None of the associations survived Benjamini-Hochberg FDR correction across phenotypes. Sample sizes were *n* = 39 for ADOS symptom dimensions and *n* = 41 for FBB-ADHD inattention. ADOS, Autism Diagnostic Observation Schedule; FBB-ADHD, parent-rated ADHD symptom questionnaire; ITPC, inter-trial phase clustering; ISPC, inter-site phase coherence.

## Discussion

In this study, we characterized how visuospatial attention is implemented across local and network-level oscillatory dynamics in autistic children and youth. We observed altered local activity during visuospatial attention in autism, accompanied by reduced reconfiguration of local and network-level dynamics when attentional demands required disengagement and reorienting.

### Task-related behavioral and neural modulation

Across groups, behavioral and neural measures differed between invalid and valid conditions. In line with established findings on visuospatial attention paradigms, we observed the expected RT cost of invalid relative to valid cues [56], and alpha power lateralization in the valid condition [13, 14, 57]. Attenuated posterior power lateralization, higher frontal beta ITPC, and higher posterior-frontal alpha phase-based connectivity in the invalid relative to the valid condition suggested greater temporal consistency and distributed coordination during attentional disengagement and reorienting across groups.

### Altered oscillatory implementation of attention in autism

Autistic participants showed an altered spatial organization of local oscillatory activity across validity conditions. In the alpha band, this was characterized by a more frontal distribution and reduced posterior lateralization, whereas neurotypical participants showed a trend towards posterior predominance, replicating findings of typical posterior alpha modulation during visuospatial attention [12, 13]. Given the proposed role of posterior alpha in selectively gating sensory input [15, 58–60], our findings may reflect a less spatially specialized implementation of attentional gating in autism. Results are in line with previous findings of a weaker posterior alpha decrease during visual and auditory processing in autistic children [31, 61–63].

Autistic participants also showed an attenuated posterior-frontal differentiation of beta power, primarily due to increased frontal beta activity, reflecting less pronounced task-related beta suppression relative to neurotypical participants. Beta oscillations have been associated with maintaining current neural states [18, 64], and the reactivation of endogenous representations [19]. Attenuated frontal beta suppression may therefore reflect stronger stabilization of internally established attentional states in autism. In addition, reduced posterior beta ITPC suggested a less consistent phase alignment of stimulus-related responses. Together, these findings point to differences in both attentional gating and maintaining attentional states in autism, which may affect altered visuospatial orienting [3] and disengagement [4].

### Reduced reconfiguration to changing attentional demands

Crucially, autistic participants presented reduced adaptation when attentional demands required disengagement and reorienting. Autistic participants showed a greater RT cost for invalid relative to valid cues, accompanied by a decreased validity-dependent modulation of neural responses. In contrast to neurotypical participants, autistic youth did not show a difference in ipsilateral alpha power between validity conditions. Attenuated adaptation to changing sensory input has also been observed for auditory and multisensory paradigms in autism [32, 42], suggesting that attenuated adaptative processing may constitute a general neural mechanism across sensory modalities.

Reduced validity condition-dependent modulation was also evident at the network level. In contrast to neurotypical participants, autistic participants showed no significant increase in posterior-frontal beta phase-based connectivity (ISPC) in the invalid relative to the valid condition, which was accompanied by a higher beta ISPC compared to neurotypical participants during the valid condition. These findings suggest stronger long-range connectivity when targets are relatively predictable, alongside reduced modulation as attentional demands change. Together with previous reports of increased task-related connectivity in autism [40–42], the findings further indicate that the direction of connectivity alterations may depend on task context.

Results of exploratory analyses on neural covariation further suggest reduced validity-dependent modulation of interactions between local and network-level measures in autism. Together, findings converge on reduced flexibility in neural responses and network dynamics as attentional demands change, consistent with previous reports of attenuated modulation of neural activity in response to increasing cognitive demands in autism [40, 43]. Flexible recruitment and coordination of brain networks are thought to support efficient allocation of neural resources by increasing integration as cognitive demands rise [65, 66]. Such demand-sensitive recruitment may itself depend on the progressive developmental specialization of functional brain systems, as frontal recruitment, long-range connectivity, and their modulation by cognitive demands change across maturation [67–72]. Accordingly, elevated connectivity to relatively predictable targets together with an attenuated modulation across neural measures may reflect less demand-sensitive regulation of network dynamics in autism, potentially relating to atypical functional specialization during brain development.

### Associations with task performance and clinical symptoms

Across groups, RT was associated with several local oscillatory measures, including measures of power, lateralization, and ITPC, consistent with previous work linking alpha and beta dynamics to response speed [12, 20, 73, 74]. Several of these neural measures also differed between groups, indicating that altered oscillatory dynamics may be relevant to visuomotor processing speed in autism [75].

Exploratory analyses within the autism group further suggested a possible link between beta dynamics, including frontal beta power, ISPC and ITPC, and the severity of restricted and repetitive behavior. Although these associations did not survive correction for multiple comparisons, their convergence in the beta band is notable given its proposed role in maintaining cognitive states [18], which may relate to behavioral inflexibility. Yet, prior work has linked beta dynamics to both repetitive and social features, while repetitive behavior has also been associated with oscillatory activity in other frequency bands [31, 40, 76, 77]. Thus, the specificity of these associations remain unclear, highlighting the need for further investigation.

## Limitations

Several limitations should be considered when interpreting the findings. First, invalidly cued targets engage multiple processes, including disengagement, reorienting, and inhibition of previously prepared motor responses. Thus, valid-invalid contrasts do not allow these components to be examined in isolation. Moreover, because analyses were baseline-corrected to the pre-cue period, cue-related preparatory activity may have carried into the target interval. Accordingly, the observed effects should be interpreted as target processing within an established attentional state rather than purely target-evoked activity. In addition, sensor-level ISPC is susceptible to volume conduction effects, although group differences and condition-dependent modulation may be less susceptible to these effects than absolute connectivity estimates.

Findings may also depend on sample characteristics, such as co-occurring ADHD symptoms. Sensitivity analyses excluding participants with elevated ADHD symptoms supported the main findings. Moreover, autism-associated behavioral alterations in visuospatial attention paradigms are not found in ADHD [78–81] and neural alterations during attentional paradigms differ between diagnoses [61, 82], suggesting that findings are unlikely to be attributable to ADHD. Additionally, groups differed in sex distribution, which was included as a covariate. Yet, replicating the effects contrasting different clinical groups, such as individuals with ADHD, with more precisely matched samples is essential.

## Conclusion

In summary, autistic children and youth showed altered local oscillatory organization across attentional contexts, together with reduced modulation of local and network-level neural dynamics when attentional demands shifted from processing expected targets towards disengagement and reorienting. Rather than reflecting a single difference in attentional fuctioning, findings point to concurrent alterations in attentional gating, maintenance of attentional states, and their adaptive reconfiguration as demands change.

## Acknowledgments

We thank Heiko Zerlaut for his support in data management and all participating families.

## Data Availability

Data is available from the corresponding author upon reasonable request. Preprocessing and analysis scripts are available at: https://github.com/LeoniePolzer/visuospatial-attention-autism-eeg.

