## Supplementary Material for "Local and network-level dynamics of alpha and beta rhythms during visuospatial attention in autism"

---

**Abbreviations:** Reaction time (RT), hemisphere (hemi), region of Interest (ROI), inter-trial phase clustering (ITPC), inter-site phase coherence (ISPC), attention-deficit/hyperactivity disorder (ADHD)

### Time-frequency representations of oscillatory power

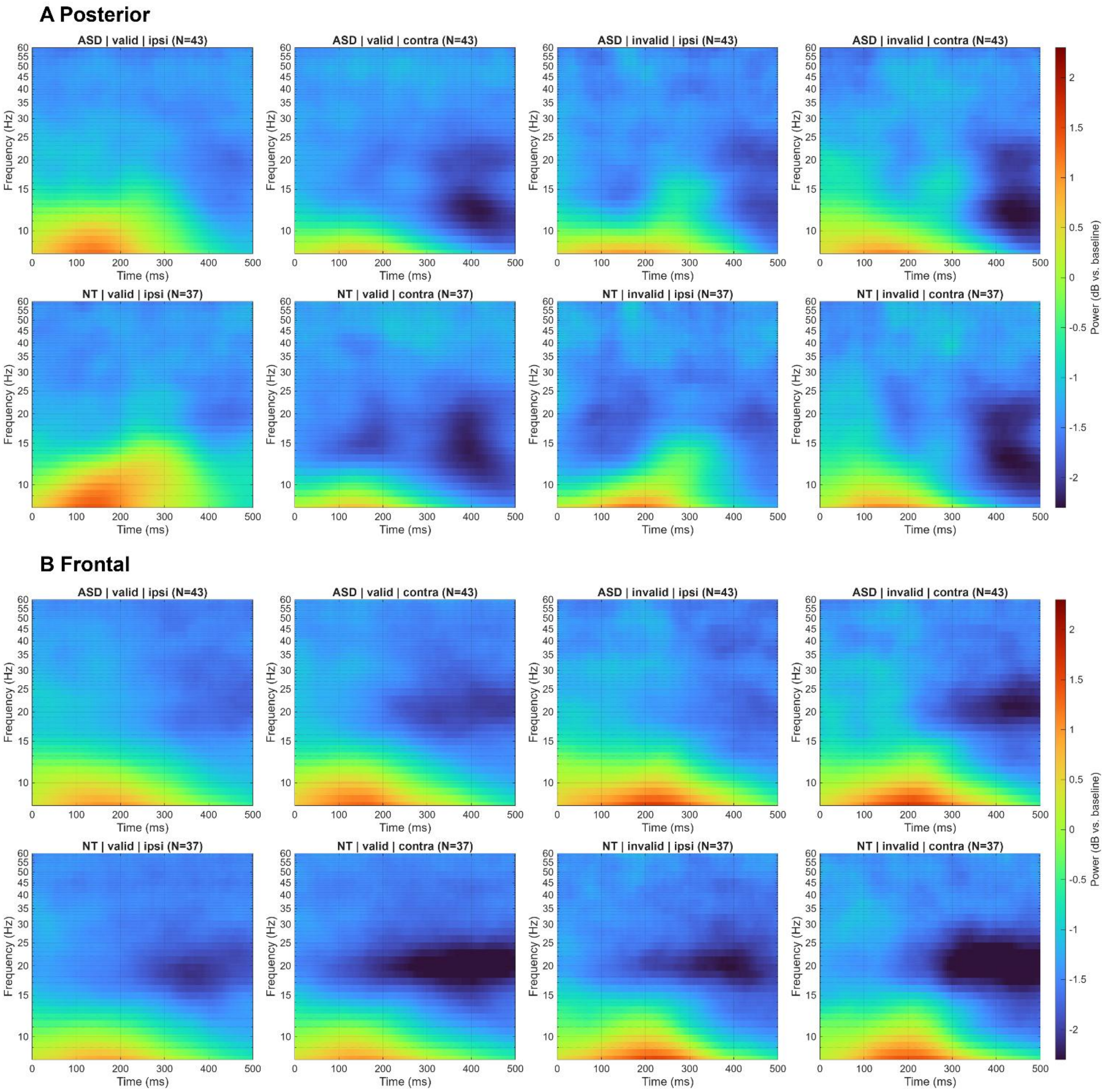

Figure S1. Target-locked time-frequency representations of oscillatory power. **Averaged time-frequency representations of baseline-corrected oscillatory power (dB) in (A) posterior and (B) frontal regions of interest (ROIs), shown separately for autistic (ASD) and neurotypical (NT) participants, valid and invalid cueing conditions, and hemispheres ipsilateral and contralateral to target location. Time zero (0 ms) indicates target onset. Warmer colors indicate increased and cooler colors decreased power relative to baseline.**

### Group differences

#### Behavioral performance

Table S1. Omnibus test: Group difference model on trial-wise RT

| Effect | F(df1, df2) | p |
|---|---|---|
| Group | 0.38 (1, 75.08) | .539 |
| Validity | 2097.53 (1, 31819.47) | < .001 |
| Age | 9.92 (1, 75.00) | .002 |
| IQ | 2.78 (1, 75.01) | .100 |
| Sex | 0.69 (1, 74.78) | .410 |
| Group × validity | 8.03 (1, 31819.55) | .005 |

#### Total power

Table S2. Omnibus test: Group difference models on trial-wise power

| Band | Hemi | Effect | F(df1, df2) | p | $p_{adj}$ |
|---|---|---|---|---|---|
| Alpha | Ipsi | Group | 0.05 (1, 75.71) | .828 | .828 |
| Alpha | Ipsi | Validity | 34.56 (1, 63720.16) | < .001 | < .001 |
| Alpha | Ipsi | ROI | 5.13 (1, 63716.05) | .023 | .023 |
| Alpha | Ipsi | Age | 0.00 (1, 75.41) | .987 | .987 |
| Alpha | Ipsi | IQ | 1.10 (1, 75.46) | .298 | .744 |
| Alpha | Ipsi | Sex | 0.59 (1, 74.62) | .444 | .713 |
| Alpha | Ipsi | Group × validity | 7.95 (1, 63720.72) | .005 | .019 |
| Alpha | Ipsi | Group × ROI | 26.55 (1, 63716.05) | < .001 | < .001 |
| Alpha | Ipsi | Validity × ROI | 100.76 (1, 63716.05) | < .001 | < .001 |
| Alpha | Ipsi | Group × validity × ROI | 1.49 (1, 63716.05) | .222 | .490 |
| Alpha | Contra | Group | 0.63 (1, 75.49) | .431 | .574 |
| Alpha | Contra | Validity | 34.71 (1, 63718.98) | < .001 | < .001 |
| Alpha | Contra | ROI | 253.06 (1, 63716.01) | < .001 | < .001 |
| Alpha | Contra | Age | 0.28 (1, 75.28) | .599 | .799 |
| Alpha | Contra | IQ | 0.39 (1, 75.31) | .534 | .744 |
| Alpha | Contra | Sex | 0.45 (1, 74.71) | .505 | .713 |
| Alpha | Contra | Group × validity | 0.34 (1, 63719.38) | .557 | .727 |
| Alpha | Contra | Group × ROI | 2.15 (1, 63716.01) | .143 | .143 |
| Alpha | Contra | Validity × ROI | 7.79 (1, 63716.01) | .005 | .007 |
| Alpha | Contra | Group × validity × ROI | 1.35 (1, 63716.01) | .245 | .490 |
| Beta | Ipsi | Group | 2.13 (1, 76.50) | .148 | .297 |
| Beta | Ipsi | Validity | 31.22 (1, 63725.95) | < .001 | < .001 |

| Band | Hemi | Effect | F(df1, df2) | p | $p_{adj}$ |
|---|---|---|---|---|---|
| Beta | Ipsi | ROI | 33.20 (1, 63715.84) | < .001 | < .001 |
| Beta | Ipsi | Age | 9.67 (1, 75.72) | .003 | .011 |
| Beta | Ipsi | IQ | 0.11 (1, 75.83) | .739 | .744 |
| Beta | Ipsi | Sex | 0.00 (1, 73.76) | .945 | .945 |
| Beta | Ipsi | Group × validity | 3.12 (1, 63727.26) | .078 | .155 |
| Beta | Ipsi | Group × ROI | 21.11 (1, 63715.84) | < .001 | < .001 |
| Beta | Ipsi | Validity × ROI | 22.14 (1, 63715.84) | < .001 | < .001 |
| Beta | Ipsi | Group × validity × ROI | 0.11 (1, 63715.84) | .740 | .740 |
| Beta | Contra | Group | 3.26 (1, 76.17) | .075 | .297 |
| Beta | Contra | Validity | 43.05 (1, 63723.74) | < .001 | < .001 |
| Beta | Contra | ROI | 42.26 (1, 63715.90) | < .001 | < .001 |
| Beta | Contra | Age | 4.73 (1, 75.59) | .033 | .066 |
| Beta | Contra | IQ | 0.11 (1, 75.67) | .744 | .744 |
| Beta | Contra | Sex | 0.39 (1, 74.07) | .535 | .713 |
| Beta | Contra | Group × validity | 0.12 (1, 63724.77) | .727 | .727 |
| Beta | Contra | Group × ROI | 11.29 (1, 63715.90) | < .001 | .001 |
| Beta | Contra | Validity × ROI | 4.80 (1, 63715.90) | .029 | .029 |
| Beta | Contra | Group × validity × ROI | 0.35 (1, 63715.90) | .557 | .740 |

#### Supplementary Information S1: Validity x ROI interactions

Validity-related modulation of power differed between frontal and posterior ROIs in all four frequency-band and hemisphere models. For ipsilateral alpha power, valid trials were associated with lower frontal power than invalid trials (b = −0.035 SD, 95% CI [−0.059, −0.012], $p_{adj}$ = .003), but higher posterior power (b = 0.135 SD, 95% CI [0.111, 0.158], $p_{adj}$ < .001). The validity effect was therefore significantly more positive in the posterior than frontal ROI (b = 0.170 SD, 95% CI [0.137, 0.203], $p_{adj}$ < .001).

For contralateral alpha power, valid trials were associated with lower power in both the frontal (b = −0.026 SD, 95% CI [−0.049, −0.003], $p_{adj}$ = .028) and posterior ROI (b = −0.073 SD, 95% CI [−0.096, −0.050], $p_{adj}$ < .001). This reduction was significantly larger posteriorly than frontally (b = −0.047 SD, 95% CI [−0.080, −0.014], $p_{adj}$ = .007).

For ipsilateral beta power, no validity effect emerged in the frontal ROI (b = 0.008 SD, 95% CI [−0.016, 0.031], $p_{adj}$ = .532), whereas posterior power was higher in valid than invalid trials (b = 0.088 SD, 95% CI [0.064, 0.112], $p_{adj}$ < .001). The posterior validity effect was significantly larger than the frontal effect (b = 0.081 SD, 95% CI [0.047, 0.114], $p_{adj}$ < .001).

For contralateral beta power, valid trials were associated with lower power in both the frontal (b = −0.037 SD, 95% CI [−0.061, −0.014], $p_{adj}$ = .002) and posterior ROI (b = −0.075 SD, 95% CI [−0.098, −0.051], $p_{adj}$ < .001). The reduction was significantly larger in the posterior ROI (b = −0.037 SD, 95% CI [−0.071, −0.004], $p_{adj}$ = .029). None of the ROI-dependent validity effects differed significantly between diagnostic groups (all group-by-validity-by-ROI $p_{adj}$ ≥ .490).

#### Lateralization

Table S3. Omnibus test: Group difference models on power lateralization

| Band | Effect | F(df1, df2) | p | $p_{adj}$ |
|---|---|---|---|---|
| Alpha | Group | 3.02 (1, 79.81) | .086 | .172 |
| Alpha | Validity | 113.16 (1, 63742.20) | < .001 | < .001 |
| Alpha | ROI | 148.62 (1, 63716.13) | < .001 | < .001 |
| Alpha | Age | 2.64 (1, 77.33) | .108 | .108 |
| Alpha | IQ | 0.56 (1, 77.65) | .455 | .883 |
| Alpha | Sex | 0.00 (1, 72.04) | .992 | .992 |
| Alpha | Group × validity | 4.27 (1, 63745.17) | .039 | .063 |
| Alpha | Group × ROI | 11.43 (1, 63716.13) | < .001 | .001 |
| Alpha | Validity × ROI | 135.60 (1, 63716.13) | < .001 | < .001 |
| Alpha | Group × validity × ROI | 0.00 (1, 63716.13) | .945 | .945 |
| Beta | Group | 1.72 (1, 89.19) | .194 | .194 |
| Beta | Validity | 109.31 (1, 63771.50) | < .001 | < .001 |
| Beta | ROI | 0.33 (1, 63717.63) | .568 | .568 |
| Beta | Age | 2.70 (1, 80.70) | .104 | .108 |
| Beta | IQ | 0.02 (1, 81.70) | .883 | .883 |
| Beta | Sex | 2.79 (1, 68.43) | .099 | .198 |
| Beta | Group × validity | 3.47 (1, 63775.60) | .063 | .063 |
| Beta | Group × ROI | 1.23 (1, 63717.63) | .267 | .267 |
| Beta | Validity × ROI | 35.58 (1, 63717.63) | < .001 | < .001 |
| Beta | Group × validity × ROI | 0.05 (1, 63717.63) | .829 | .945 |

##### Supplementary Information S2: Post-hoc comparisons on lateralization effects

For alpha lateralization, significant group-by-ROI (F(1, 63,716.13) = 11.43, $p_{adj}$ = .001), and validity-by-ROI interactions emerged (F(1, 63,716.13) = 135.60, $p_{adj}$ < .001). Valid relative to invalid trials were associated with increased lateralization in the posterior ROI (b = 0.191 SD, 95% CI [0.167, 0.215], $p_{adj}$ < .001), but not in the frontal ROI (b = −0.009 SD, 95% CI [−0.032, 0.015], $p_{adj}$ = .638). The validity effect was significantly larger posteriorly than frontally (b = 0.200 SD, 95% CI [0.166, 0.233], $p_{adj}$ < .001). Accordingly, posterior and frontal lateralization did not differ in invalid trials ($p_{adj}$ = .743), whereas posterior lateralization was higher in valid trials (b = 0.204 SD, 95% CI [0.186, 0.223], $p_{adj}$ < .001).

Both groups showed greater posterior than frontal alpha lateralization, but this regional difference was smaller in autistic participants (ASD: b = 0.075 SD, 95% CI [0.052, 0.099], $p_{adj}$ < .001; NT: b = 0.133 SD, 95% CI [0.109, 0.158], $p_{adj}$ < .001; between-group difference: b = −0.058 SD, 95% CI [−0.092, −0.024], $p_{adj}$ = .001). Neurotypical participants additionally showed higher posterior alpha lateralization than autistic participants (b = 0.059 SD, 95% CI [0.021, 0.096], $p_{adj}$ = .003), whereas no group difference emerged frontally ($p_{adj}$ = .976).

For beta lateralization, a significant validity-by-ROI interaction was observed (F(1, 63,717.63) = 35.58, $p_{adj}$ < .001). Valid relative to invalid trials increased lateralization in both the frontal (b = 0.039 SD, 95% CI [0.015, 0.063], $p_{adj}$ = .001) and posterior ROI (b = 0.141 SD, 95% CI [0.118, 0.165], $p_{adj}$ < .001), with a significantly larger increase posteriorly (b = 0.103 SD, 95% CI [0.069, 0.136], $p_{adj}$ < .001). Posterior

lateralization was lower than frontal lateralization in invalid trials (b = −0.056 SD, 95% CI [−0.084, −0.028], $p_{adj} < .001$), but higher in valid trials (b = 0.046 SD, 95% CI [0.028, 0.065], $p_{adj} < .001$).
Group-by-validity interactions did not survive correction for multiple comparisons in either frequency band (both $p_{adj}$ = .063), and neither group-by-validity-by-ROI interaction was significant (both $p_{adj} \geq$ .945).

#### Inter-trial phase clustering (ITPC)

Table S4. Omnibus test: Group difference models on ITPC

| Band | Effect | F(df1, df2) | p | $p_{adj}$ |
|---|---|---|---|---|
| Alpha | Group | 0.18 (1, 74.84) | .670 | .670 |
| Alpha | Validity | 0.15 (1, 232.55) | .704 | .704 |
| Alpha | ROI | 112.58 (1, 231.91) | < .001 | < .001 |
| Alpha | Age | 2.39 (1, 74.72) | .126 | .252 |
| Alpha | IQ | 0.09 (1, 74.91) | .762 | .835 |
| Alpha | Sex | 3.19 (1, 74.80) | .078 | .157 |
| Alpha | Group × validity | 0.67 (1, 232.55) | .415 | .415 |
| Alpha | Group × ROI | 0.00 (1, 231.91) | .952 | .952 |
| Alpha | Validity × ROI | 0.75 (1, 231.91) | .386 | .386 |
| Alpha | Group × validity × ROI | 0.02 (1, 231.91) | .895 | .895 |
| Beta | Group | 1.57 (1, 75.20) | .213 | .427 |
| Beta | Validity | 0.65 (1, 233.48) | .422 | .704 |
| Beta | ROI | 176.40 (1, 232.37) | < .001 | < .001 |
| Beta | Age | 0.42 (1, 75.01) | .518 | .518 |
| Beta | IQ | 0.04 (1, 75.32) | .835 | .835 |
| Beta | Sex | 0.68 (1, 75.13) | .412 | .412 |
| Beta | Group × validity | 2.48 (1, 233.48) | .116 | .233 |
| Beta | Group × ROI | 5.66 (1, 232.37) | .018 | .036 |
| Beta | Validity × ROI | 5.65 (1, 232.37) | .018 | .037 |
| Beta | Group × validity × ROI | 3.39 (1, 232.37) | .067 | .134 |

##### Supplementary Information S3: Validity x ROI interactions

Validity-related modulation of ITPC differed significantly between frontal and posterior ROIs in the beta band (F(1, 232) = 5.65, $p_{adj}$ = .037), but not in the alpha band (F(1, 232) = 0.75, $p_{adj}$ = .386). For beta ITPC, valid trials were associated with lower frontal ITPC than invalid trials (b = −0.238 SD, 95% CI [−0.446, −0.031], $p_{adj}$ = .033), whereas no significant validity effect emerged in the posterior ROI (b = 0.118 SD, 95% CI [−0.090, 0.325], $p_{adj}$ = .267). The validity effect was significantly more positive in the posterior than frontal ROI (b = 0.356 SD, 95% CI [0.062, 0.649], $p_{adj}$ = .037). This ROI-dependent validity effect did not significantly differ between diagnostic groups (Group × Validity × ROI: $p_{adj}$ = .134).

#### Power-based connectivity

Table S5. Omnibus test: Group difference models on power connectivity

| Band | Effect | F(df1, df2) | p | $p_{adj}$ |
|---|---|---|---|---|
| Alpha | Group | 2.28 (1, 75.00) | .135 | .270 |
| Alpha | Validity | 0.36 (1, 78.00) | .549 | .773 |
| Alpha | Age | 7.14 (1, 75.00) | .009 | .009 |
| Alpha | IQ | 1.43 (1, 75.00) | .236 | .471 |
| Alpha | Sex | 0.77 (1, 75.00) | .382 | .382 |
| Alpha | Group × validity | 0.83 (1, 78.00) | .364 | .728 |
| Beta | Group | 0.13 (1, 75.00) | .722 | .722 |
| Beta | Validity | 0.08 (1, 78.00) | .773 | .773 |
| Beta | Age | 19.12 (1, 75.00) | < .001 | < .001 |
| Beta | IQ | 0.29 (1, 75.00) | .592 | .592 |
| Beta | Sex | 1.04 (1, 75.00) | .310 | .382 |
| Beta | Group × validity | 0.00 (1, 78.00) | .987 | .987 |

#### Inter-site phase coherence (ISPC)

Table S6. Omnibus test: Group difference models on inter-site phase coherence (ISPC)

| Band | Effect | F(df1, df2) | p | $p_{adj}$ |
|---|---|---|---|---|
| Alpha | Group | 0.25 (1, 75.09) | .617 | .617 |
| Alpha | Validity | 5.60 (1, 77.28) | .020 | .020 |
| Alpha | Age | 14.04 (1, 75.01) | < .001 | < .001 |
| Alpha | IQ | 0.31 (1, 75.14) | .578 | .669 |
| Alpha | Sex | 0.09 (1, 75.06) | .765 | .765 |
| Alpha | Group × validity | 3.76 (1, 77.28) | .056 | .056 |
| Beta | Group | 7.16 (1, 74.94) | .009 | .018 |
| Beta | Validity | 10.08 (1, 77.07) | .002 | .004 |
| Beta | Age | 24.69 (1, 74.88) | < .001 | < .001 |
| Beta | IQ | 0.18 (1, 74.97) | .669 | .669 |
| Beta | Sex | 0.90 (1, 74.92) | .347 | .693 |
| Beta | Group × validity | 7.25 (1, 77.07) | .009 | .017 |

### Supplementary Information S4: Simulation-based power analyses

To evaluate the sensitivity of the study design to the planned interaction effects, we conducted Monte Carlo simulation-based power analyses using mixed-effects models matching the structure of the primary analyses. Simulations were based on the observed sample sizes and repeated-measures structure after preprocessing. Outcome values were generated under predefined standardized interaction effects. Total outcome variance was standardized to 1, with ICC = .50 used as the primary planning assumption and ICCs of .30 and .70 examined as sensitivity conditions. Power was estimated using Type III tests with Satterthwaite degrees of freedom.

For the trial-level power analyses, sensitivity was high even for small standardized interaction effects. Under the primary ICC = .50 assumption, a group-by-validity interaction of 0.05 was detected with 98.8% power, and power for the group-by-validity-by-ROI interaction increased from 56.6% at an effect of 0.05 to 98.8% at 0.10. Participant-level analyses also showed good sensitivity to moderate two-way interactions: approximately 80% power was reached for standardized group-by-validity effects of about 0.45 for ITPC and 0.65 for connectivity. As expected for the more complex higher-order contrast, sensitivity was lower for the participant-level group-by-validity-by-ROI ITPC interaction, for which an effect of approximately 0.90 was required to reach 80% power under the primary ICC assumption. The overall pattern was consistent across ICC assumptions of .30–.70, with particularly high sensitivity in the trial-level analyses and good sensitivity to moderate two-way interaction effects in the participant-level models.

#### Sensitivity analyses: Exclusion of participants with elevated ADHD symptoms

Four participants scored above the cut-off for ADHD in a clinical ADHD questionnaire (FBB-ADHD; Döpfner et al., 2008). We re-analyzed models that showed group differences in neural measures to test whether these group differences were potentially driven by co-occuring ADHD. The main effects of interest remained qualitatively unchanged after the exclusion of these participants.

Table S7. Omnibus tests of group-difference models for power after exclusion of participants with elevated ADHD symptoms.

| Band | Hemi | Effect | F(df1, df2) | p | $p_{adj}$ |
|---|---|---|---|---|---|
| Alpha | Ipsi | Group | 0.09 (1, 71.69) | .765 | .765 |
| Alpha | Ipsi | Validity | 31.16 (1, 61047.69) | < .001 | < .001 |
| Alpha | Ipsi | ROI | 2.68 (1, 61044.03) | .102 | .102 |
| Alpha | Ipsi | Age | 0.23 (1, 71.27) | .631 | .841 |
| Alpha | Ipsi | IQ | 0.59 (1, 71.38) | .445 | .814 |
| Alpha | Ipsi | Sex | 0.61 (1, 70.69) | .438 | .665 |
| Alpha | Ipsi | Group × validity | 8.60 (1, 61048.15) | .003 | .013 |
| Alpha | Ipsi | Group × ROI | 19.92 (1, 61044.03) | < .001 | < .001 |
| Alpha | Ipsi | Validity × ROI | 104.82 (1, 61044.03) | < .001 | < .001 |
| Alpha | Ipsi | Group × validity × ROI | 0.61 (1, 61044.03) | .435 | .870 |
| Alpha | Contra | Group | 0.67 (1, 71.46) | .416 | .555 |
| Alpha | Contra | Validity | 34.21 (1, 61046.61) | < .001 | < .001 |
| Alpha | Contra | ROI | 245.62 (1, 61044.00) | < .001 | < .001 |
| Alpha | Contra | Age | 0.01 (1, 71.17) | .933 | .933 |
| Alpha | Contra | IQ | 0.12 (1, 71.25) | .735 | .814 |
| Alpha | Contra | Sex | 0.46 (1, 70.75) | .499 | .665 |
| Alpha | Contra | Group × validity | 0.43 (1, 61046.94) | .512 | .683 |
| Alpha | Contra | Group × ROI | 2.35 (1, 61044.00) | .126 | .126 |
| Alpha | Contra | Validity × ROI | 6.93 (1, 61044.00) | .008 | .011 |
| Alpha | Contra | Group × validity × ROI | 1.08 (1, 61044.00) | .298 | .870 |
| Beta | Ipsi | Group | 2.44 (1, 72.62) | .123 | .245 |
| Beta | Ipsi | Validity | 28.37 (1, 61053.22) | < .001 | < .001 |
| Beta | Ipsi | ROI | 39.43 (1, 61043.92) | < .001 | < .001 |
| Beta | Ipsi | Age | 13.26 (1, 71.49) | < .001 | .002 |
| Beta | Ipsi | IQ | 0.06 (1, 71.78) | .814 | .814 |
| Beta | Ipsi | Sex | 0.02 (1, 70.01) | .897 | .897 |
| Beta | Ipsi | Group × validity | 3.51 (1, 61054.33) | .061 | .122 |
| Beta | Ipsi | Group × ROI | 14.91 (1, 61043.92) | < .001 | < .001 |

| Band | Hemi | Effect | F(df1, df2) | p | $p_{adj}$ |
|---|---|---|---|---|---|
| Beta | Ipsi | Validity × ROI | 24.13 (1, 61043.92) | < .001 | < .001 |
| Beta | Ipsi | Group × validity × ROI | 0.00 (1, 61043.92) | .984 | .984 |
| Beta | Contra | Group | 3.71 (1, 72.31) | .058 | .232 |
| Beta | Contra | Validity | 45.15 (1, 61051.32) | < .001 | < .001 |
| Beta | Contra | ROI | 45.72 (1, 61043.97) | < .001 | < .001 |
| Beta | Contra | Age | 8.24 (1, 71.43) | .005 | .011 |
| Beta | Contra | IQ | 0.20 (1, 71.66) | .660 | .814 |
| Beta | Contra | Sex | 0.51 (1, 70.26) | .479 | .665 |
| Beta | Contra | Group × validity | 0.00 (1, 61052.21) | .957 | .957 |
| Beta | Contra | Group × ROI | 8.43 (1, 61043.97) | .004 | .005 |
| Beta | Contra | Validity × ROI | 3.97 (1, 61043.97) | .046 | .046 |
| Beta | Contra | Group × validity × ROI | 0.18 (1, 61043.97) | .674 | .898 |

Table S8. Omnibus tests of group-difference models for lateralization after exclusion of participants with elevated ADHD symptoms.

| Band | Effect | F(df1, df2) | p | $p_{adj}$ |
|---|---|---|---|---|
| Alpha | Group | 2.59 (1, 75.46) | .112 | .179 |
| Alpha | Validity | 106.95 (1, 61067.12) | < .001 | < .001 |
| Alpha | ROI | 157.85 (1, 61043.86) | < .001 | < .001 |
| Alpha | Age | 2.25 (1, 72.17) | .138 | .258 |
| Alpha | IQ | 0.73 (1, 72.89) | .394 | .431 |
| Alpha | Sex | 0.00 (1, 68.41) | .986 | .986 |
| Alpha | Group × validity | 4.44 (1, 61069.63) | .035 | .070 |
| Alpha | Group × ROI | 7.27 (1, 61043.86) | .007 | .014 |
| Alpha | Validity × ROI | 136.89 (1, 61043.86) | < .001 | < .001 |
| Alpha | Group × validity × ROI | 0.05 (1, 61043.86) | .826 | .826 |
| Beta | Group | 1.84 (1, 85.64) | .179 | .179 |
| Beta | Validity | 107.55 (1, 61096.24) | < .001 | < .001 |
| Beta | ROI | 0.12 (1, 61045.48) | .734 | .734 |
| Beta | Age | 1.30 (1, 74.47) | .258 | .258 |
| Beta | IQ | 0.63 (1, 76.30) | .431 | .431 |
| Beta | Sex | 3.06 (1, 65.51) | .085 | .170 |
| Beta | Group × validity | 2.88 (1, 61099.99) | .090 | .090 |
| Beta | Group × ROI | 0.75 (1, 61045.48) | .386 | .386 |
| Beta | Validity × ROI | 35.71 (1, 61045.48) | < .001 | < .001 |
| Beta | Group × validity × ROI | 0.12 (1, 61045.48) | .731 | .826 |

Table S9. Omnibus tests of group-difference models for ITPC after exclusion of participants with elevated ADHD symptoms.

| Band | Effect | F(df1, df2) | p | $p_{adj}$ |
|---|---|---|---|---|
| Alpha | Group | 0.02 (1, 70.78) | .896 | .896 |
| Alpha | Validity | 0.32 (1, 220.49) | .575 | .575 |
| Alpha | ROI | 108.68 (1, 219.88) | < .001 | < .001 |
| Alpha | Age | 0.61 (1, 70.70) | .439 | .681 |
| Alpha | IQ | 0.00 (1, 70.90) | .949 | .949 |
| Alpha | Sex | 3.57 (1, 70.75) | .063 | .126 |
| Alpha | Group × validity | 0.98 (1, 220.49) | .323 | .323 |
| Alpha | Group × ROI | 0.01 (1, 219.88) | .931 | .931 |
| Alpha | Validity × ROI | 0.83 (1, 219.88) | .364 | .364 |
| Alpha | Group × validity × ROI | 0.00 (1, 219.88) | .952 | .952 |
| Beta | Group | 1.01 (1, 71.17) | .319 | .638 |
| Beta | Validity | 0.47 (1, 221.40) | .494 | .575 |
| Beta | ROI | 161.33 (1, 220.38) | < .001 | < .001 |
| Beta | Age | 0.17 (1, 71.03) | .681 | .681 |
| Beta | IQ | 0.01 (1, 71.38) | .904 | .949 |
| Beta | Sex | 0.79 (1, 71.11) | .378 | .378 |
| Beta | Group × validity | 2.05 (1, 221.40) | .154 | .308 |
| Beta | Group × ROI | 5.76 (1, 220.38) | .017 | .034 |
| Beta | Validity × ROI | 6.35 (1, 220.38) | .012 | .025 |
| Beta | Group × validity × ROI | 2.39 (1, 220.38) | .123 | .246 |

Table S10. Omnibus tests of group-difference models for power connectivity after exclusion of participants with elevated ADHD symptoms.

| Band | Effect | F(df1, df2) | p | $p_{adj}$ |
|---|---|---|---|---|
| Alpha | Group | 2.43 (1, 71.00) | .124 | .248 |
| Alpha | Validity | 0.14 (1, 74.00) | .709 | .931 |
| Alpha | Age | 4.42 (1, 71.00) | .039 | .039 |
| Alpha | IQ | 2.01 (1, 71.00) | .161 | .321 |
| Alpha | Sex | 0.76 (1, 71.00) | .386 | .386 |
| Alpha | Group × validity | 1.22 (1, 74.00) | .273 | .546 |
| Beta | Group | 0.02 (1, 71.00) | .899 | .899 |
| Beta | Validity | 0.01 (1, 74.00) | .931 | .931 |
| Beta | Age | 21.23 (1, 71.00) | < .001 | < .001 |
| Beta | IQ | 0.98 (1, 71.00) | .326 | .326 |
| Beta | Sex | 1.31 (1, 71.00) | .257 | .386 |

| Band | Effect | F(df1, df2) | p | $p_{adj}$ |
|---|---|---|---|---|
| Beta | Group × validity | 0.06 (1, 74.00) | .815 | .815 |

Table S11. Omnibus tests of group-difference models for ISPC after exclusion of participants with elevated ADHD symptoms.

| Band | Effect | F(df1, df2) | p | $p_{adj}$ |
|---|---|---|---|---|
| Alpha | Group | 0.25 (1, 71.08) | .620 | .620 |
| Alpha | Validity | 6.15 (1, 73.26) | .015 | .015 |
| Alpha | Age | 8.72 (1, 71.02) | .004 | .004 |
| Alpha | IQ | 0.63 (1, 71.15) | .431 | .511 |
| Alpha | Sex | 0.10 (1, 71.06) | .755 | .755 |
| Alpha | Group × validity | 3.10 (1, 73.26) | .083 | .083 |
| Beta | Group | 6.36 (1, 70.94) | .014 | .028 |
| Beta | Validity | 12.84 (1, 73.07) | < .001 | .001 |
| Beta | Age | 21.88 (1, 70.91) | < .001 | < .001 |
| Beta | IQ | 0.44 (1, 71.00) | .511 | .511 |
| Beta | Sex | 0.97 (1, 70.93) | .328 | .656 |
| Beta | Group × validity | 5.70 (1, 73.07) | .020 | .039 |

#### Behavioral performance prediction by neural measures

Table S12. Omnibus test: Trial-wise power effects on reaction time (RT)

| Band | Hemisphere | Effect | F(df1, df2) | p | $p_{adj}$ |
|---|---|---|---|---|---|
| Alpha | Contralateral | Validity | 2108.81 (1, 31816.54) | < .001 | < .001 |
| Alpha | Contralateral | Posterior power | 0.61 (1, 31837.49) | .437 | .437 |
| Alpha | Contralateral | Frontal power | 3.94 (1, 31826.61) | .047 | .094 |
| Alpha | Contralateral | Age | 11.06 (1, 75.99) | .001 | .001 |
| Alpha | Contralateral | IQ | 3.50 (1, 76.03) | .065 | .066 |
| Alpha | Contralateral | Sex | 0.97 (1, 75.77) | .327 | .327 |
| Alpha | Contralateral | Validity × posterior power | 0.17 (1, 31818.57) | .679 | .906 |
| Alpha | Contralateral | Validity × frontal power | 0.03 (1, 31817.72) | .871 | .893 |
| Alpha | Ipsilateral | Validity | 2058.54 (1, 31816.61) | < .001 | < .001 |
| Alpha | Ipsilateral | Posterior power | 14.54 (1, 31832.14) | < .001 | < .001 |
| Alpha | Ipsilateral | Frontal power | 3.94 (1, 31826.87) | .047 | .094 |
| Alpha | Ipsilateral | Age | 11.12 (1, 75.99) | .001 | .001 |
| Alpha | Ipsilateral | IQ | 3.49 (1, 76.04) | .066 | .066 |
| Alpha | Ipsilateral | Sex | 0.98 (1, 75.78) | .325 | .327 |
| Alpha | Ipsilateral | Validity × posterior power | 9.89 (1, 31820.05) | .002 | .007 |
| Alpha | Ipsilateral | Validity × frontal power | 2.78 (1, 31818.19) | .096 | .383 |
| Beta | Contralateral | Validity | 2114.49 (1, 31816.52) | < .001 | < .001 |
| Beta | Contralateral | Posterior power | 4.65 (1, 31824.49) | .031 | .042 |
| Beta | Contralateral | Frontal power | 1.37 (1, 31824.36) | .242 | .242 |
| Beta | Contralateral | Age | 11.08 (1, 76.00) | .001 | .001 |
| Beta | Contralateral | IQ | 3.51 (1, 76.04) | .065 | .066 |
| Beta | Contralateral | Sex | 0.98 (1, 75.78) | .325 | .327 |
| Beta | Contralateral | Validity × posterior power | 1.25 (1, 31817.62) | .264 | .528 |
| Beta | Contralateral | Validity × frontal power | 0.15 (1, 31817.49) | .696 | .893 |
| Beta | Ipsilateral | Validity | 2094.71 (1, 31816.54) | < .001 | < .001 |
| Beta | Ipsilateral | Posterior power | 6.81 (1, 31824.59) | .009 | .018 |
| Beta | Ipsilateral | Frontal power | 2.89 (1, 31824.32) | .089 | .119 |
| Beta | Ipsilateral | Age | 11.26 (1, 76.00) | .001 | .001 |
| Beta | Ipsilateral | IQ | 3.51 (1, 76.04) | .065 | .066 |
| Beta | Ipsilateral | Sex | 0.99 (1, 75.78) | .322 | .327 |
| Beta | Ipsilateral | Validity × posterior power | 0.01 (1, 31818.63) | .917 | .917 |
| Beta | Ipsilateral | Validity × frontal power | 0.02 (1, 31818.22) | .893 | .893 |

Table S13. Omnibus test: Lateralization effects on reaction time (RT)

| Band | Effect | F(df1, df2) | p | $p_{adj}$ |
|---|---|---|---|---|
| Alpha | Validity | 2055.74 (1, 31816.56) | < .001 | < .001 |
| Alpha | Posterior lateralization | 10.22 (1, 31821.57) | .001 | .003 |
| Alpha | Frontal lateralization | 3.13 (1, 31817.93) | .077 | .077 |
| Alpha | Age | 11.11 (1, 76.00) | .001 | .001 |
| Alpha | IQ | 3.48 (1, 76.04) | .066 | .066 |
| Alpha | Sex | 0.98 (1, 75.78) | .325 | .326 |
| Alpha | Validity × posterior lateralization | 3.77 (1, 31820.49) | .052 | .105 |
| Alpha | Validity × frontal lateralization | 0.16 (1, 31817.74) | .687 | .895 |
| Beta | Validity | 2082.92 (1, 31816.54) | < .001 | < .001 |
| Beta | Posterior lateralization | 2.24 (1, 31817.77) | .135 | .135 |
| Beta | Frontal lateralization | 8.48 (1, 31817.50) | .004 | .007 |
| Beta | Age | 11.11 (1, 76.00) | .001 | .001 |
| Beta | IQ | 3.50 (1, 76.04) | .065 | .066 |
| Beta | Sex | 0.98 (1, 75.78) | .326 | .326 |
| Beta | Validity × posterior lateralization | 0.77 (1, 31818.12) | .380 | .380 |
| Beta | Validity × frontal lateralization | 0.02 (1, 31817.58) | .895 | .895 |

Table S14. Omnibus test: ITPC effects on reaction time (RT)

| Band | Effect | F(df1, df2) | p | $p_{adj}$ |
|---|---|---|---|---|
| Alpha | Validity | 484.32 (1, 66.45) | < .001 | < .001 |
| Alpha | Posterior ITPC | 3.96 (1, 94.51) | .050 | .099 |
| Alpha | Frontal ITPC | 6.41 (1, 92.59) | .013 | .026 |
| Alpha | Age | 11.19 (1, 68.08) | .001 | .001 |
| Alpha | IQ | 3.71 (1, 67.35) | .058 | .060 |
| Alpha | Sex | 2.83 (1, 68.08) | .097 | .194 |
| Alpha | Validity × posterior ITPC | 0.17 (1, 72.05) | .681 | .681 |
| Alpha | Validity × frontal ITPC | 1.02 (1, 72.86) | .316 | .316 |
| Beta | Validity | 459.49 (1, 70.86) | < .001 | < .001 |
| Beta | Posterior ITPC | 2.24 (1, 83.72) | .138 | .138 |
| Beta | Frontal ITPC | 0.03 (1, 84.36) | .855 | .855 |
| Beta | Age | 11.73 (1, 71.10) | .001 | .001 |
| Beta | IQ | 3.67 (1, 71.32) | .060 | .060 |
| Beta | Sex | 1.26 (1, 70.86) | .265 | .265 |
| Beta | Validity × posterior ITPC | 1.08 (1, 74.05) | .301 | .602 |
| Beta | Validity × frontal ITPC | 4.76 (1, 74.32) | .032 | .065 |

Table S15. Omnibus test: Power connectivity effects on reaction time (RT)

| Band | Effect | F(df1, df2) | p | $p_{adj}$ |
|---|---|---|---|---|
| Alpha | Validity | 457.79 (1, 77.18) | < .001 | < .001 |
| Alpha | Power connectivity | 0.01 (1, 92.19) | .925 | .925 |
| Alpha | Age | 11.26 (1, 77.46) | .001 | .001 |
| Alpha | IQ | 3.48 (1, 76.06) | .066 | .066 |
| Alpha | Sex | 1.07 (1, 75.94) | .304 | .304 |
| Alpha | Validity × power connectivity | 0.03 (1, 81.27) | .860 | .860 |
| Beta | Validity | 479.69 (1, 77.03) | < .001 | < .001 |
| Beta | Power connectivity | 3.08 (1, 97.81) | .082 | .165 |
| Beta | Age | 13.29 (1, 79.84) | < .001 | < .001 |
| Beta | IQ | 3.71 (1, 75.80) | .058 | .066 |
| Beta | Sex | 1.24 (1, 75.95) | .269 | .304 |
| Beta | Validity × power connectivity | 2.40 (1, 79.91) | .126 | .251 |

Table S16. Omnibus test: ISPC effects on reaction time (RT)

| Band | Effect | F(df1, df2) | p | $p_{adj}$ |
|---|---|---|---|---|
| Alpha | Validity | 479.03 (1, 78.61) | < .001 | < .001 |
| Alpha | Phase connectivity | 2.36 (1, 143.92) | .127 | .253 |
| Alpha | Age | 8.39 (1, 81.96) | .005 | .005 |
| Alpha | IQ | 3.61 (1, 75.99) | .061 | .075 |
| Alpha | Sex | 1.26 (1, 75.69) | .265 | .297 |
| Alpha | Validity × phase connectivity | 0.57 (1, 76.95) | .454 | .454 |
| Beta | Validity | 460.92 (1, 80.60) | < .001 | < .001 |
| Beta | Phase connectivity | 0.00 (1, 151.46) | .997 | .997 |
| Beta | Age | 10.11 (1, 84.40) | .002 | .004 |
| Beta | IQ | 3.26 (1, 75.72) | .075 | .075 |
| Beta | Sex | 1.10 (1, 75.59) | .297 | .297 |
| Beta | Validity × phase connectivity | 1.40 (1, 77.16) | .240 | .454 |
